# Zinc-embedded fabrics inactivate SARS-CoV-2 and influenza A virus

**DOI:** 10.1101/2020.11.02.365833

**Authors:** Vikram Gopal, Benjamin E. Nilsson-Payant, Hollie French, Jurre Y. Siegers, Wai-shing Yung, Matthew Hardwick, Aartjan J.W. te Velthuis

**Affiliations:** Ascend Performance Materials, 1010 Travis Street, Suite 900, Houston, TX 77002, USA; Department of Microbiology, Icahn School of Medicine at Mount Sinai, New York, NY 10029, USA; Division of Virology, Department of Pathology, Addenbrooke’s Hospital, University of Cambridge, Hills Road, CB2 2QQ, United Kingdom; Department of Viroscience, Erasmus University Medical Centre, Rotterdam, the Netherlands; ResInnova Laboratories, 8807 Colesville Rd, 3rd Floor, Silver Spring, MD 20910, USA

**Keywords:** influenza, coronavirus, absorption, zinc, face mask

## Abstract

Infections with respiratory viruses can spread via liquid droplets and aerosols, and cause diseases such as influenza and COVID-19. Face masks and other personal protective equipment (PPE) can act as barriers that prevent the spread of respiratory droplets containing these viruses. However, influenza A viruses and coronaviruses are stable for hours on various materials, which makes frequent and correct disposal of these PPE important. Metal ions embedded into PPE may inactivate respiratory viruses, but confounding factors such as absorption of viruses make measuring and optimizing the inactivation characteristics difficult. Here we used polyamide 6.6 (PA66) fibers that had zinc ions embedded during the polymerisation process and systematically investigated if these fibers can absorb and inactivate pandemic SARS-CoV-2 and influenza A virus H1N1. We find that these viruses are readily absorbed by PA66 fabrics and inactivated by zinc ions embedded into this fabric. The inactivation rate (pfu·gram^−1^·min^−1^) exceeds the number of active virus particles expelled by a cough and supports a wide range of viral loads. Moreover, we found that the zinc content and the virus inactivating property of the fabric remain stable over 50 standardized washes. Overall, these results provide new insight into the development of “pathogen-free” PPE and better protection against RNA virus spread.

## Introduction

Infections with influenza A viruses (IAV), influenza B viruses (IBV) and coronaviruses (CoV) are a burden on our healthcare systems and economy. These respiratory RNA viruses transmit through aerosols, liquid droplets and fomites and seasonal strains typically cause a mild disease with symptoms including nasopharyngitis, fever, coughing, and headache. Nevertheless, seasonal IAV and IBV result in 290,000-645,000 deaths every year and billions of dollars in losses, in part due to increased hospitalizations and reduced work efficiencies.

The impact of highly pathogenic and pandemic IAV and CoV strains is even more severe. Over the past century, several pandemic influenza A virus (IAV) and severe acute respiratory syndrome coronaviruses (SARS-CoV) strains have infected and killed millions of people. Of particular importance were the 1918 H1N1, 1957 H2N2, and the 1968 H3N2 pandemic IAV strains, and the SARS-CoV-2 pandemic strain, the causative agent of COVID-19. These viruses can cause viral pneumonia and make it easier for secondary bacterial infections to take hold, increasing patient morbidity and mortality (1). Understanding how we can efficiently prevent the spread of these viruses will be important for current and future RNA virus outbreaks.

IAV is part of the *Orthomyxoviridae*, a family of negative-sense RNA viruses that are known for their segmented RNA genomes. The virus particle is enveloped by a double-layered membrane and contains multiple copies of the viral haemagglutinin (HA), matrix 2 (M2), and, neuraminidase (NA) proteins embedded in the membrane (2). The HA protein binds sialic acid receptors on the outside of host cells, and fuses the viral membrane and the cellular plasma membrane, while the M2 protein acts as proton channel that plays a role in the activation of HA and release of the viral RNA genome into the host cell. The viral RNA genome consists of eight segments that are encapsidated by the viral nucleoprotein (NP) and RNA polymerase as ribonucleoprotein (RNP) complexes inside virus particles (3). After viral transcription, protein synthesis, replication and virion formation, the NA protein is required for the release of virus particles from the host cell.

SARS-CoV-2 belongs to the *Coronaviridae*, a family of positive-sense RNA viruses that infect a wide range of vertebrates, including humans (4, 5). The SARS-CoV-2 virion consists of a double-layered membrane and membrane proteins spike (S), envelope (E) and matrix (M). The viral RNA genome is harboured inside the virus particle and encapsidated by the viral nucleocapsid protein (N). Infection of a host cell requires binding of the S protein to the cellular receptor ACE2 (6). Following entry, the virus releases its viral RNA into the host cell for viral protein synthesis and genome transcription and replication by the viral RNA polymerase.

Various antivirals are currently available for the treatment of influenza and CoV infections, including RNA polymerase inhibitors favipiravir and remdesivir (7–9). In addition, vaccines are available or in development that target the HA or S protein to prevent infection and spread of IAV and SARS-CoV-2 (10). Unfortunately, vaccines are not readily available for emerging RNA viruses, and existing RNA viruses can become resistant against antivirals and escape immune pressures due to their relatively high mutation rate (11, 12). Use of personal protective equipment (PPE), such as face masks, is therefore recommended by health organizations to prevent respiratory virus spread and several studies have supported their efficacy (13–15). However, opponents of the use of face masks have pointed to complicating factors, including observations that respiratory viruses are stable for days to hours on fabrics and that N95 respirators require careful decontamination to allow their reuse (16–19). Further considerations are the additional environmental waste that disposable face masks produce, the poor fit of some masks, and the potential health hazard that discarded masks present (20, 21). Development of PPE that can trap and inactivate respiratory viruses may help address some of these concerns or simplify their use in day-to-day life.

Previous research has shown that IAVs and CoVs can be inactivated by metal surfaces, such as copper and zinc (22–24). While the exact underlying inactivation mechanisms are not fully understood, evidence suggests that metal ions can induce RNA hydrolysis, membrane destabilisation, or viral protein inactivation or degradation (25–27). So far, few studies have investigated if metal ions embedded in fabrics can inactivate RNA viruses, in part because absorbance and fabric density differences among fabrics present confounding factors that the protocols approved for testing the inactivating properties of surfaces do not account for.

To tackle some of these confounding issues, we here measured the ability of different fabrics, such as cotton, polyamide 6.6 (PA66) and polypropylene (PPP), to trap H1N1 IAV and pandemic SARS-CoV-2, and we explore how we can remove these viruses from the fabrics to test for inactivation. We find that cotton and PA66 readily absorb respiratory viruses, and that zinc ions embedded in PA66-based fabrics resulted in approximately a 2-log reduction in virus titer, which is more than sufficient to inactivate the number of infectious IAV virus particles (~24 plaque forming units [pfu]) present in a cough (28). Virus inactivation plateaued over time. Overall, these results provide new insight into the protective properties of fabrics used for face masks and the development of “pathogen-free” fabrics.

## Results

### Influenza virus absorbance by cotton, polypropylene and polyamide

Various studies have investigated the filtration properties of fabrics and have investigated factors such as breathability, hydrophobicity and/or electrostaticity. Fabrics also have a different weight per square meter (gram/m^2^) and different moisture retention abilities, with cotton absorbing up to 500% of its weight and PA66 absorbing as little as 0.3% of its weight, depending on environmental conditions (29). These different properties may affect how fabrics trap and/or release aerosols or liquid droplets containing RNA viruses. For instance, PPE with poor absorbing properties that become contaminated with viruses may retain these viruses on their surface thus become a potential health hazard if not disposed of properly. Presently, it is not fully understood how moisture retention is correlated with virus particle absorption. To investigate this relationship, we added IAV strain A/WSN/33 (H1N1) to International Antimicrobial Council (IAC) issued cotton, a textile PA66 fabric, or PPP from a disposable type II 3-ply face mask (Fig. 1A). After a 30-min incubation at room temperature, the fabrics were washed with PBS to remove unabsorbed virus (Fig. 1B). To estimate the amount of remaining liquid on each fabric, each sample tube with fabric was weighed and compared to its dry weight. As shown in Fig. 1C and D, cotton and PA66 retained more liquid than PPP, both relative to the applied volume and the weight of the fabric. Subsequent analysis of the IAV titer in the input and fabric washes showed that the cotton and woven PA66 fabrics readily absorbed the applied virus, while less virus was observed by the PPP fabric (Fig. 1E, F), which is in line with the hydrophobicity of PPP (30).

**Figure 1.**
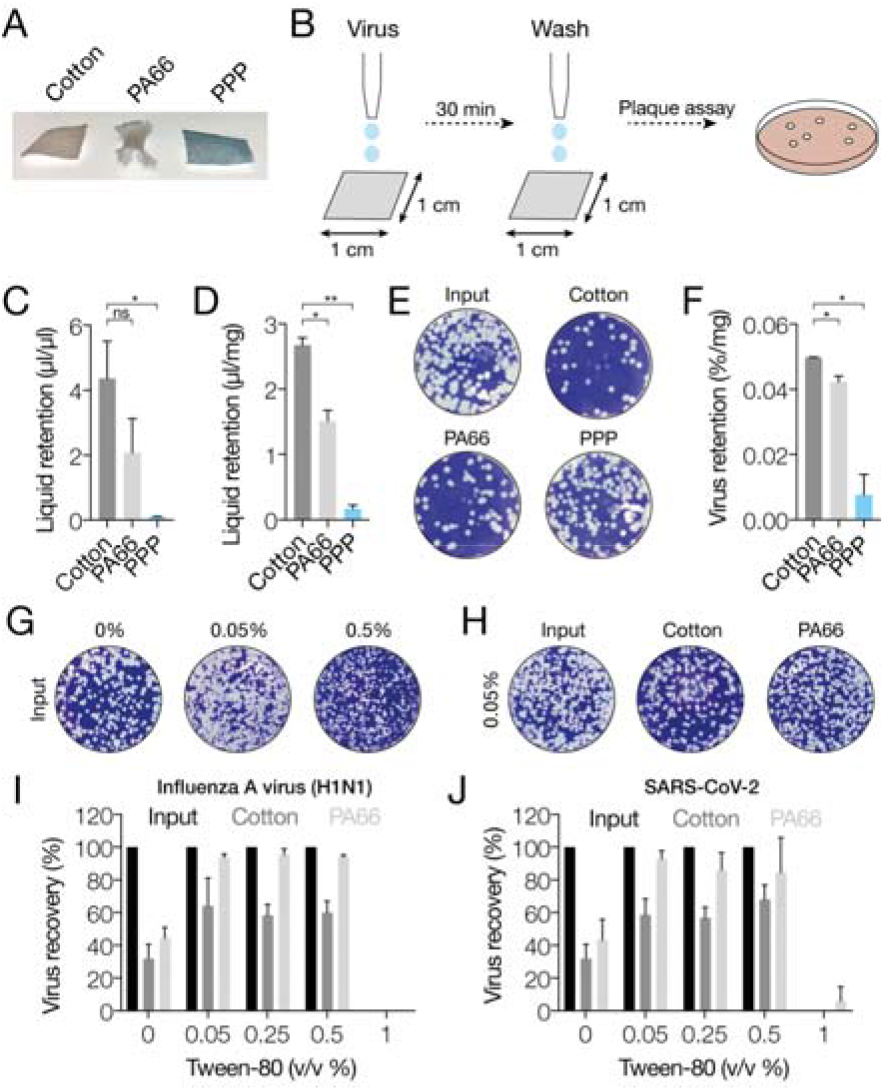
Absorption and release of IAV and SARS-CoV-2 from fabrics. **A**) Photographs of cotton control, PA66 and polypropylene fabric samples. **B**) Schematic of experimental procedure for exposing and isolating RNA virus from fabrics. **C**) Analysis of virus medium retention by fabrics per volume of input medium. Values were obtained by weighing each fabric before and after addition of virus medium, and after removal of the virus medium. **D**) Analysis of virus medium retention by fabrics normalized by dry weight of each fabric. Values were obtained by weighing each fabric before and after addition of virus medium, and after removal of the virus medium. **E**) Plaque assay of IAV present in virus medium after removal of the medium from each fabric. **F**) Quantitation of the amount of virus remaining on each fabric, normalized by the dry weight of each fabric. **G**) Effect of different tween-80 concentrations on IAV plaque assay read-out. **H**) Effect of 0.05% tween-80 in PBS on the amount of virus released from each fabric. **I**) Quantitation of IAV titers after absorption of the virus to the fabrics and washing of the fabrics with PBS or PBS containing different concentrations of tween-80. **J**) Quantitation of SARS-CoV-2 titers after absorption of the virus to the fabrics and washing of the fabrics with PBS or PBS containing different concentrations of tween-80. Error bars indicate standard deviation. Asterisk indicates p-value, with * p<0.05, ** p<0.005, and ns p>0.05.

In order to remove IAV from the cotton and PA66 fabrics without inactivating the virus, we added different concentrations of polysorbate-80 (tween-80) - a mild detergent that is also used in IAV vaccine preparations - to the PBS wash buffer (Fig. 1G). We did not observe any cytopathic effects of the detergent on the Madin-Darby Canine Kidney (MDCK) cells used for the plaque assay, but did find that the presence of 0.05%-0.1% tween-80 increased the apparent viral titer relative to infections in PBS (Fig. 1G), whereas 0.25-0.5% tween-80 reduced the apparent IAV plaque size (Fig. 1G). We found that 0.05% tween-80 succeeded in recovering more than 94% of the virus from the PA66 woven fabric, whereas 61% was removed from the cotton fabric (Fig. 1H and I). Higher concentrations, such as 1% tween-80, prevented IAV infection (Fig. 1I).

To confirm whether other viruses can be removed from cotton and woven PA66 as well, we repeated the experiment with SARS-CoV-2. We found that over 92% of SARS-CoV-2 can be recovered from the woven PA66 fabric using 0.05% tween-80, while up to 59% could be recovered from the cotton fabric (Fig. 1J). Together, these results demonstrate that IAV and SARS-CoV-2 are strongly absorbed by cotton and PA66, suggesting that these materials would trap respiratory viruses inside face masks. At the same time, these findings imply that PPP is poor at trapping respiratory viruses. Since IAV and SARS-CoV-2 can be removed from a PA66 fabric with a mild detergent, this protocol can be useful for testing the inactivating properties of fabrics.

### Influenza virus is inactivated by zinc ions

Copper and zinc surfaces or particles can inactivate IAV strains and seasonal CoV HCoV-229E, and PPP imbued with copper oxide can potentially inactivate IAV (22, 25, 26, 31). For embedding into polymers, zinc ions provide benefits over copper ions as zinc has a much higher propensity to ionize than copper, and thereby provides a much faster reaction potential. Moreover, zinc oxide, which we embedded in the PA66 polymer used here, is considered a Generally regarded as Safe (GRAS) compound by the FDA, which can speed up the development process. Finally, zinc does not cause discolouration of the polymer or fabric enabling a broader applicability. However, like copper, zinc ions are cytotoxic in tissue culture (Fig. 2A), which confounds analysis of their effect on viral titers. We found that addition of an equimolar concentration of EDTA following the virus incubation with zinc ions (Fig. 2B) can efficiently chelate zinc ions and prevent cytotoxic effects (Fig. 2C). EDTA alone does not have any cytotoxic effects and does not reduce viral titers (Fig. 2A).

**Figure 2.**
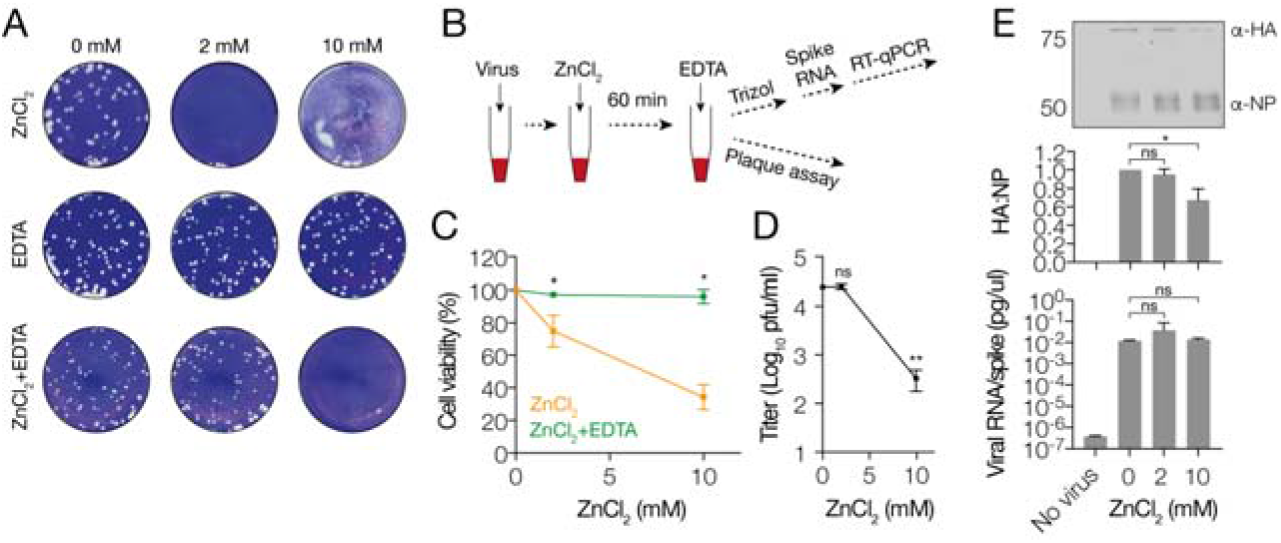
IAV is inactivated by zinc ions. **A**) Plaque assay showing the effect of different zinc chloride and EDTA concentrations on IAV titers. **B**) Experimental approach for inactivating IAV with zinc ions and neutralization of zinc ions using EDTA. **C**) Cytotoxicity analysis of zinc chloride and EDTA in MDCK cells. **D**) IAV titers after exposure to zinc chloride and neutralization with EDTA as measured on MDCK cells. **E**) Western blot IAV HA and NP protein levels after exposure to zinc chloride and neutralization with EDTA. Upper panel shows quantitation of western signal and middle panel the western signal as detected with LI-COR. Bottom panel shows NA segment RT-qPCR analysis of IAV virus after exposure to zinc chloride and neutralization with EDTA. Error bars represent standard deviation. Asterisk indicates p-value, with * p<0.05, ** p<0.005, and ns p>0.05.

To investigate if zinc ions can directly inactivate IAV, we incubated influenza virus with varying concentrations of zinc chloride. After 60 min, the reactions were stopped with an equimolar amount of EDTA and subsequently diluted for virus titer determination by plaque assay (Fig. 2B). As shown in Fig. 2D, we found that addition of zinc chloride resulted in a significant reduction in the IAV titer. Previous research has shown that metal ions can destabilize viral proteins (25). To gain more insight into the mechanism of virus inactivation, viral protein levels in the zinc chloride-treated samples were analysed by western blot. As shown in Fig. 2E, we found that in the presence of zinc chloride, HA levels were reduced in a concentration-dependent manner, while NP levels did not diminish (Fig. 2E). This result thus suggest that zinc ions may affect the IAV surface proteins more significantly than the internal proteins. To test if IAV RNA levels were affected, we added a 120-nucleotide long spike RNA to each sample, extracted viral RNA, and performed reverse transcriptions (RT) using a 31 terminal NA primer. cDNA levels were next quantified using quantitative polymerase chain reaction (qPCR) of the NA gene-encoding segment and normalized to the spike RNA level (Fig. 2E). No effect of zinc chloride on viral NA segment levels was found. Together, these results imply that zinc ions can inactivate an IAV (H1N1) strain by destabilization of the viral surface proteins.

### Influenza and coronavirus strains are inactivated on fabrics containing zinc ions

The above results suggest that zinc ions can directly inactivate an IAV H1N1 strain. To investigate if these inactivating properties are also present when zinc ions are embedded in a PA66 matrix, we used 0.4 gram of a textile PA66 fabric containing 328 ppm zinc ions (equivalent to 2.5 mM; abbreviated as KF1). Incubation of KF1 with virus and washing of the fabrics using a PBS buffer containing 0.05% tween-80 and 10 mM EDTA (PBSTE; Fig. 3A), resulted in an approximately 2-log reduction of the IAV and SARS-CoV-2 titers compared to a PA66 control fabric after 1 h (Fig. 3B and C). To confirm that inactivation of these viruses occurred on KF1, viral protein levels were analysed in the PBSTE wash eluate by western blot (Fig. 3D and E). Any virus that remained in the fabric after extraction with PBSTE, was lysed and extracted using Trizol and analyzed by western blot as well. Western blots showed a reduction in the HA and S protein level in the virus eluate that was removed from the KF1 fabric compared to the control fabric eluate for IAV and SARS-CoV-2, respectively (Fig. 3D and E). The signal obtained from the virus that remained on each fabric after the PBSTE extraction was close to background, in line with the observations in Fig. 1, and we were only able to quantify the SARS-CoV-2 signal, but observed no statistically significant difference. Overall, we conclude that inactivation of IAV and SARS-CoV-2 occurs on a fabric embedded with zinc oxide, analogous to the previously observed effects of copper oxide (31). To better investigate the rate of reduction, we incubated KF1 with virus for different lengths of time and subtracted the absorbed virus titer in the negative control from the level of reduction in the KF1 fabric and fitted the data with a logarithmic equation (Fig. 3F). A maximum reduction occurred between 30 seconds and 5 min of incubation, and the virus titer reduction reached a plateau after approximately 50 min.

**Figure 3.**
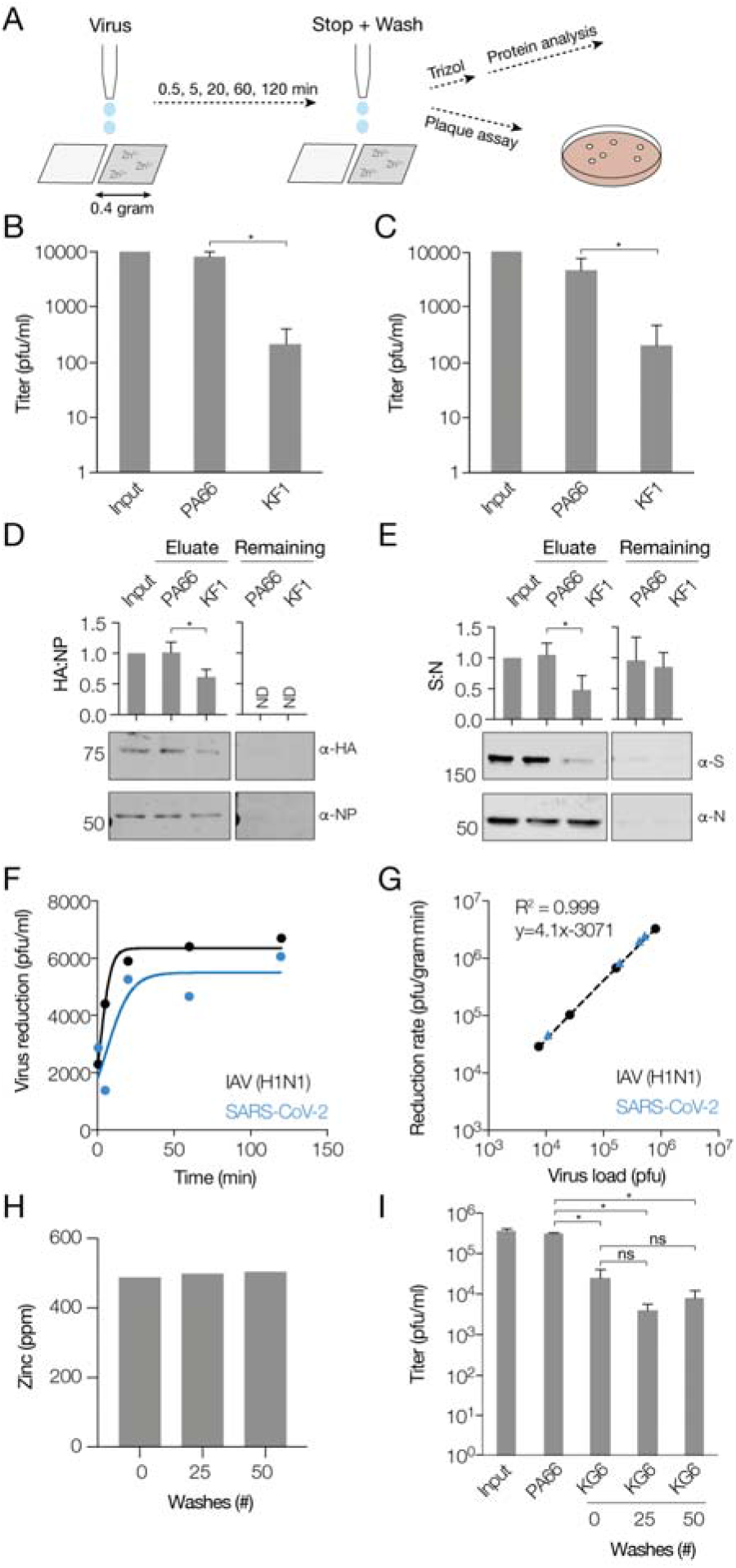
Inactivation of IAV and SARS-CoV-2 on fabrics. **A**) Schematic of testing procedure for fabrics without or with embedded zinc oxide. **B**) IAV titer in input, or PA66 control or KF1 fabric eluates. **C**) SARS-CoV-2 titer in input, or PA66 control or KF1 fabric eluates. One representative experiment is shown. **D**) Western blot analysis of IAV HA and NP protein levels after exposure of IAV to the KF1 or control fabric. Both the virus that was removed (eluate) from each fabric with PBSTE as well as the virus that remained on each fabric was analyzed. **E**) Western blot analysis of SARS-CoV-2 S and N protein levels after exposure of virus to the KF1 or control fabric. Both the virus that was removed (eluate) from each fabric with PBSTE as well as the virus that remained on each fabric was analyzed. **F**) Time course of IAV or SARS-CoV-2 titer reduction by the KF1 fabric minus the titer reduction by the PA66 control without embedded zinc. One representative time course is shown. Data were fit with logarithmic equation. **G**) Reduction rate of IAV or SARS-CoV-2 titer after exposure to KF1 fabric. Data points were obtained by from time courses experiments in which we varied the viral load and subsequently estimated the maximum reduction rate (exponential phase) for each time course. Reduction was normalized to pfu·gram^−1^·min^−1^ using the dry fabric weight. IAV and SARS-CoV-2 data points were fit with a linear line and no difference was observed between the two fits. R^2^ for IAV fit is shown. **H**) Zinc content in KG6 fabric after repeated washing according to the standardized home laundry test protocol AATCC M6-2016. One representative experiment for one batch of fabric is shown. **I**) Reduction rate of IAV titer after exposure to unwashed or washed KG6 fabric. Error bars represent standard deviation. Asterisk indicates p-value, with * p<0.05 and ns p>0.05.

### Inactivation of IAV and coronaviruses scales with virus load

To investigate the robustness and saturation level of the inactivation by fabrics containing embedded zinc oxide, we next performed experiments with KF1 and varied the viral load added to each fabric over a range of 10^3^ to 10^7^ pfu. The liquid volume applied to each fabric was kept constant. After incubation for different periods of time, fabrics were washed with PBSTE, virus titers estimated by plaque assay and the virus titer reduction rate calculated based on the shortest incubation time. Reduction rates were subsequently normalized by the dry weight of each fabric. As shown in Fig. 3G, the rate of reduction in virus titer (in pfu·gram^−1^·min^−1^) scaled with virus load. On a log-log plot, the data could be fit with a linear equation. To confirm the robustness of these findings, we performed the same experiments with SARS-CoV-2, and found a similar behavior (Fig. 3H).

To investigate if fabrics constructed from fibers containing zinc ions maintain their zinc oxide content after washing, a KF1 fabric with 500 ppm zinc ions (equivalent to 5.3 mM; internal code KG6) was washed 25 or 50 times using the standardized home laundry test protocol AATCC M6-2016. Subsequent analysis of the zinc content after washing revealed that the zinc content remained relatively constant in the PA66 fabrics for up to 50 washes (Fig. 3H). We next confirmed if these washed fabrics were still able to reduce virus titers and incubated 0.4 g of unwashed or washed fabric with a fixed amount of IAV and removed inactivated virus with PBSTE. Analysis of the virus titers showed that both washed fabrics were able reduce the IAV titer by approximately 2-logs (Fig. 3I). Overall, these results suggest that the PA66 fabric containing zinc can inactivate both IAV and SARS-CoV-2 and that this property is retained after 50 washes.

## Discussion

Infections with respiratory RNA viruses cause regular seasonal epidemics and occasional pandemics, and thus present a severe burden on our personal health, healthcare systems, and economy. While seasonal respiratory viruses - including over 160 different rhinoviruses, human CoVs strains NL63, OC43, HKU1 and 229E, influenza A, B and C viruses, human respiratory syncytial virus, human parainfluenza viruses, and human metapneumovirus – typically cause mild disease, IAV and CoVs have also been associated with zoonotic outbreaks and lethal pandemics. Vaccines and antivirals are available or in development for various respiratory viruses, but the appearance of resistance to antivirals or vaccines is a known or potential problem. In the absence of new vaccines or antivirals, one way to fight RNA viruses is to limit respiratory virus spread through efficient PPE.

To better understand how respiratory RNA viruses are absorbed and inactivated on fabrics, we here added IAV and SARS-CoV-2 to cotton, PA66, and PPP. We find strong absorption by cotton and PA66 in the standard laboratory buffer PBS, and that addition of tween-80 results in efficient virus release from PA66, but not from cotton. A previous clinical trial found that cotton masks with strong absorbing properties may be associated with a higher risk of infection when reused and our finding that cotton does not release IAV or SARS-CoV-2 efficiently after washing is in line with this observation (16). By contrast, virus retention on PPP, which is used for the construction of disposable 3-ply masks, is poor, in line with its hydrophobic properties (30). This result implies that respiratory viruses remain on the surface of these masks and together with findings that SARS-CoV-2 can survive on various surfaces for several hours to days, and even 7 days on PPP-based surgical face masks (19, 32), PPP-based masks may increase the risk of infection if not handled and disposed of properly. However, PPP has of course alternative properties, such as good breathability, filtration, and electrostatic properties and will thus has a purpose in the right situation.

The use of any face mask or other PPE, even if used temporarily but correctly to prevent spread, is better than wearing no face mask as it may reduce the risk of infection with respiratory viruses (13–15). However, the above considerations suggest that there is a potential for the use of pathogen-inactivating PPE, i.e. fabrics that can both absorb as well as inactivate viruses. In particular the use of zinc and copper ions in PPE is promising in this regard, as these metals can inactivate IAV and SARS-CoV-2 (Fig. 2, 3) (22, 24, 26). Using a PA66-based fabric from which we could easily remove absorbed virus with a mild detergent (Fig. 1), we tested the effect of zinc ions embedded in a matrix on the infectiousness of IAV and pandemic SARS-CoV-2. We consistently found a rapid reduction in the titer of all viruses tested and at viral loads that far exceed the number of infectious IAV particles present in a cough (Fig. 3). After washing the fabrics using a standardized protocol, both the zinc content as well as the inactivating properties of the PA66 fabric were retained, suggesting that this fabric is reusable at least 50 times. This property may be of particular importance for designing reusable PPE that could help reduce environmental waste, virus transmission, and costs.

We also investigated the mechanism by which zinc ions inactivate IAV and SARS-CoV-2. RT-qPCR analysis showed no significant reduction in viral RNA integrity after treatment with zinc ions. By contrast, analysis of the stability of the viral surface and capsid proteins revealed a reduced stability of the virus surface proteins HA and S, for IAV and SARS-CoV-2, respectively, after exposure to zinc ions, while no effect on the internal nucleoprotein or nucleocapsid proteins was detected. We observed a similar altered surface protein to nucleoprotein ratio after exposure to the zinc containing PA66 fabric KF1. Together, these results suggest that the reduction in virus titer after exposure to zinc ions derives from inactivation of the viral surface proteins. This is in line with previous research using copper ions (25). Research has shown that zinc and copper ions can also induce oxidative reactions, inactivation of the viral proton channels, or viral membrane destabilization and we cannot exclude that these processes may play a role in the inactivation as well (27, 33, 34).

Overall, these results strongly suggest that virus inactivating fabrics can offer enhanced safety over widely used cotton and PPP-based PPE. Our findings may therefore be important for health care workers who are exposed to infected patients for prolonged periods, people with underlying risk factors needing additional protection, and people who need to frequently remove their PPE.

## Methods

### Influenza viruses and cells

MDCK cells were originally sources from ATCC. Influenza A/WSN/33 (H1N1) virus was rescues from plasmids (35) and grown on MDCK cells in Minimal Essential Medium (MEM) containing 0.5% foetal bovine serum (FBS) at 37 °C and 5% CO_2_. Plaque assays were performed on 100% confluent MDCK cells in MEM containing 0.5% FBS with a 1% agarose overlay. Ten-fold virus dilutions were grown under a 1% agarose in MEM containing 0.5% FBS overlay for 2 days at 37 °C. Cell viability was measured using a CellTiter Blue assay (Promega).

### Coronaviruses and cells

SARS-CoV-2 (Bavpat-1 and USA-WA1/2020) were grown on African Green Monkey kidney epithelial Vero-E6 cells in Dulbecco’s Minimal Essential Medium (DMEM) supplemented with 10% FBS. For plaque assay analysis, Vero-E6 cells were seeded in 12-well plates and infected at 100% confluency. Ten-fold virus dilutions were grown under a 1% agarose overlay in DMEM containing 0.5% FBS for 2 days at 37 °C. Experiments were performed in a BSL3 lab according to approved biosafety standards.

### Fabrics

Fabric samples were cut by cleaned scissors. The cotton fabric was issued and certified by the IAC (lot number IACVC01012020). The PA66 fabrics with zinc ions (Microban Additive Zo7; EPA Reg. No. 42182-8) and a control fabric without zinc ions were produced and provided by Ascend Performance Materials. The PPP disposable type II 3-ply face mask (Medical Products Co, Ltd) was BS EN14638:2019 type II compliant. PA66 fabrics were washed according to the standardized home laundry test protocol AATCC M6-2016. Inductively coupled plasma (ICP) analysis was used to determine the zinc content after fabric washing.

### Virus absorption and extraction

To test the ability of fabrics to reduce viral titres, we used a modified ISO 18184 protocol. Briefly, 100 μl of IAV strain A/WSN/33 (H1N1) was applied in 5-10 μl droplets to cotton, a textile PA66 fabric, or PPP cut from medical grade face masks. The size of the fabrics varied from 1 cm^2^ to 0.4 gram, as indicated in the figures. After an incubation at room temperature as indicated, the fabrics were washed with PBS, PBS containing tween-80, or PBS containing 0.05% tween-80 and 10 mM EDTA through vortexing. After virus removal, 1 ml of Trizol (Invitrogen) was added to each fabric to extract remaining viral protein and RNA. Experiments were performed in triplicate, unless noted otherwise. Data was analysed in Graphpad Prism 8 using 1-way ANOVA.

### RT-qPCR and western blot

RNA extraction from Trizol was performed as described previously(36), while protein was extracted from the interphase using isopropanol precipitation(37). Precipitated protein was washed in ethanol, resuspended in 5x SDS-PAGE loading buffer, sonicated for 10 seconds, and boiled for 10 min before 8% SDS-PAGE analysis. Western blot was performed using antibodies directed against IAV HA (Invitrogen, PA5-34929) and NP (GeneTex, GTX125989) and SARS-CoV-2 S (Abcam ab272504) and N (GeneTex, GTX632269). Membranes were washed in TBS containing 0.1% tween-20. Spike RNA was purchased from IDT and had the sequence 5⍰-AGUAGAAACAAGGCGGUAGGCGCUGUCCUUUAUCCAGACAACCAUUACCUGUCCACACA AUCUGCCCUUUCGAAAGAUCCCAACGAAAAGAGAGACCACAUGGUCCUUCCUGCUUUUGCU-3⍰. Isolated RNA was reverse transcribed using SuperScript III and a primer binding to the 3’ end of the NA segment (36). qPCR was performed as described previously (36). Data was analysed in Graphpad Prism 8 using one-way ANOVA with multiple corrections.

## Acknowledgments

We thank Shanaka Rodrigo, Natasha Virjee, Chris Hsiung, and Benjamin Tenoever for helpful discussions, materials and information. AtV is supported by joint Wellcome Trust and Royal Society grant 206579/Z/17/Z and the National Institutes of Health grant R21AI147172.

